# Mortality among adults living with HIV treated for tuberculosis based on positive, negative, or no bacteriologic test results for tuberculosis: the IeDEA consortium

**DOI:** 10.1101/571000

**Authors:** John M Humphrey, Philani Mpofu, April C. Pettit, Beverly Musick, E. Jane Carter, Eugene Messou, Olivier Marcy, Brenda Crabtree-Ramirez, Marcel Yotebieng, Kathryn Anastos, Timothy R. Sterling, Constantin Yiannoutsos, Lameck Diero, Kara Wools-Kaloustian

**Affiliations:** Department of Medicine, Indiana University School of Medicine, Indianapolis, IN, United States; Department of Biostatistics, Indiana University Fairbanks School of Public Health, Indianapolis, IN, United States; Vanderbilt University Medical Center, Nashville, TN, United States; Vanderbilt Tuberculosis Center, Nashville, TN, United States; Department of Medicine, Brown University School of Medicine, Providence, RI, United States; University of Bordeaux, Centre INSERM U1219, Bordeaux Population Health, Bordeaux, France; Centre de Prise en charge de Recherche et de Formation (Aconda-CePReF), Abidjan, Côte d’Ivoire; Epidemiology and Public Health Unit, Institut Pasteur du Cambodge, Phnom Penh, Cambodia; Instituto Nacional de Ciencias Médicas y Nutrición Salvador Zubirán, Mexico City, Mexico; The Ohio State University, College of Public Health, Columbus, OH, United States; Department of Medicine, Albert Einstein College of Medicine, Bronx, NY, United States; Department of Medicine, Moi University College of Health Sciences, Eldoret, Kenya

**Author notes:** Membership in the IeDEA consortium for the participating programs is provided in the Supporting Information (S8 Table).

## Abstract

**Background:** In resource-constrained settings, people living with HIV (PLWH) treated for tuberculosis (TB) despite negative bacteriologic tests have a higher mortality than those treated with positive tests. Many PLWH are treated without bacteriologic testing; their mortality compared to those with bacteriologic testing is uncertain.

**Methods:** We conducted an observational cohort study among PLWH ≥ 15 years of age who initiated TB treatment at clinical sites affiliated with four regions of the International epidemiology Databases to Evaluate AIDS (IeDEA) consortium from 2012-2014: Caribbean, Central and South America, and Central, East, and West Africa. The primary exposure of interest was the TB bacteriologic test status at TB treatment initiation: positive, negative, or no test result. The hazard for death in the 12 months following TB treatment initiation was estimated using the Cox proportional hazard model, adjusted for patient- and site-level factors. Missing covariates were multiply imputed.

**Results:** Among 2,091 PLWH included, the median age at TB treatment initiation was 36 years, 44% were female, 53% had CD4 counts ≤ 200 cells/mm^3^, and 52% were on antiretroviral treatment (ART). Compared to patients with positive bacteriologic tests, the adjusted hazard for death was higher among patients with no test results (HR 1.56, 95% CI 1.08-2.26) but not different than those with negative tests (HR 1.28, 95% CI 0.91-1.81). Older age was also associated with a higher hazard for death, while being on ART, having a higher CD4 count, West Africa region, and tertiary facility level were associated with lower hazards for death.

**Conclusion:** PLWH treated for TB with no bacteriologic test results were more likely to die than those treated with positive tests, underscoring the importance of TB bacteriologic diagnosis in resource-constrained settings. Research is needed to understand the causes of death among PLWH treated for TB in the absence of positive bacteriologic tests.

## Introduction

Although tuberculosis (TB) accounted for 300,000 deaths among people living with HIV (PLWH) in 2017, diagnosing TB in resource-limited settings remains a challenge [1]. In 2017, only 56% of the 5.5 million pulmonary TB cases reported to the World Health Organization (WHO) globally were bacteriologically confirmed (i.e. positive for smear microscopy, culture, or nucleic acid amplification test [NAAT]) [1]. Among studies reporting the autopsy prevalence of TB in HIV-related deaths, TB was prevalent in 37% of deaths, but in half of those cases, TB was not diagnosed by the time of death [2]. The rollout of nucleic acid amplification tests (NAAT) such as the Xpert^®^ MTB/RIF (Cepheid, Sunnyvale, CA, USA), which are more sensitive and specific than smear microscopy, has helped close this diagnostic gap [3]. However, limited impact on mortality has been observed with the use of Xpert MTB/RIF, in part due to high baseline rates of empiric TB treatment (i.e. based on clinical symptoms or radiographic signs) in high TB burden settings [3–5]. A better understanding of the risk of death among PLWH in the context of varying TB test results, or the absence of TB testing, is needed to inform the management of PLWH treated for TB in resource-limited settings where the risk of TB is high.

Acquiring a bacteriologic diagnosis of TB in resource-limited settings can be hampered by economic, clinical and test-related factors. Despite recent increases in TB diagnostic test coverage in sub-Saharan Africa, smear microscopy and culture is estimated to be available at only 1.4 and 0.7 laboratories per 100,000 population, respectively [6, 7]. Many labs in resource-constrained settings suffer from weak supply systems, outdated equipment, poor quality control, and insufficient staffing [6, 8–10]. Smear microscopy for acid-fast bacilli (AFB) is often the only TB test available in such settings but it is poorly sensitive among PLWH, with 30-60% of pulmonary TB cases reported to be smear negative [11–13]. Clinicians may not order bacteriologic testing for TB among PLWH due to lack of knowledge about TB (or conversely, knowledge of the limitations of smear microscopy), or in cases of suspected extrapulmonary TB requiring invasive tissue sampling that is not feasible to perform [5, 8, 14, 15]. Patients may not be able to produce a sputum sample for bacteriologic testing or access TB testing sites (e.g. for serial sputum collection) due to the distance or cost of transport [8, 16, 17].

Empiric TB treatment in PLWH, either because a bacteriologic test was negative or no test was performed, is thus common in resource-limited settings [4, 5, 18–20]. However, mortality in the absence of TB bacteriologic testing is not well defined [21–24]. The objective of this study was to describe the characteristics and risk of death among PLWH treated for TB in the context of positive, negative, or no TB bacteriologic test results.

## Materials and Methods

### Study setting and patient population

This observational cohort study utilized data previously collected from PLWH who were enrolled in HIV care programs affiliated with four participating regions of the International epidemiology Databases to Evaluate AIDS (IeDEA) consortium: Caribbean, Central and South America (CCASAnet), and Central, East, and West Africa [25]. IeDEA is a National Institutes of Health (NIH)-funded consortium that pools and harmonizes baseline and follow-up patient data collected in the context of routine care [26]. All participating facilities provided standard of care HIV and TB treatment services per their respective national guidelines. The study population included all PLWH ≥ 15 years of age who initiated TB treatment between January 2012 and December 2014. Patients were excluded if an alternative diagnosis was established and TB therapy was stopped. Recurrent TB cases within the study period were excluded (n = 41), so a patient could not contribute more than one TB case. Patients receiving a drug-resistant TB treatment regimen were also excluded, which was defined according to WHO criteria as any injectable agent (except streptomycin), fluoroquinolones, or oral bacteriostatic agent (e.g. para-aminosalicylic acid, ethionamide, cycloserine) [27]. The reporting of this study conforms to the STROBE statement (S1 Table) [28].

### Ethics statement

Independent Ethics Committee (IEC) or Institutional Review Board (IRB) approval for this study was obtained by each of the local IeDEA sites as well as from the Indiana University IRB (S2 Table).

### Data management

TB cases were identified at participating IeDEA sites through review of local TB registries by site staff. A standardized, electronic case report form developed in Research Electronic Data Capture (REDCap) and available in English and French was used to collect medical record data [29]. Data entry into case report forms was conducted from January 2012 through January 2016 by local IeDEA investigators following medical record review. Patient-level HIV data were obtained from the IeDEA regional HIV care and treatment data repositories for all patients with completed case report forms. Site-level data were obtained from surveys of antiretroviral treatment (ART) sites participating in IeDEA conducted between March 1 and July 1, 2012 [30, 31]. Routine audits were performed to ensure data quality throughout data collection. Regional IeDEA data were transferred to the IeDEA-East Africa Regional Data Center where they were merged, and additional data quality assessments were undertaken before analysis.

### Study definitions

Adult patients were defined as individuals ≥ 15 years of age to be consistent with WHO and other national and regional TB programs [32, 33]. The primary exposure of interest was the TB bacteriologic test status at the time of TB treatment initiation: *Positive test* group included patients with one or more positive results including acid fast bacilli (AFB) smear, culture, or NAAT; *Negative test* group included patients for whom one or more of these bacteriologic tests were performed and none were positive; *No test* group includes patients for whom no bacteriologic test was performed or results reported [34]. Patient-level variables at TB treatment initiation included: age, sex, body mass index (BMI), ART status at TB treatment initiation (on ART vs. not on ART), CD4 count (defined as the nearest value within 180 days before or 30 days after TB treatment initiation), TB disease site (pulmonary vs. extrapulmonary, and specific extrapulmonary site(s)), and type of bacteriologic TB test (smear, culture, NAAT; the specimen type [e.g. respiratory vs. non-respiratory] was not available). TB disease site was categorized into pulmonary and extrapulmonary according to the WHO and CDC reporting definitions [32, 34, 35]. Accordingly, pulmonary TB included any case involving the lung parenchyma or tracheobronchial tree, and miliary TB with lesions in the lungs; extrapulmonary TB included cases involving organs other than the lungs; cases of combined pulmonary and extrapulmonary TB were classified as pulmonary TB.

Site-level variables included IeDEA region (CCASAnet, Central Africa, East Africa, West Africa), setting (urban, peri-urban [i.e. immediately adjoining urban areas], facility level (secondary [i.e district or provincial facilities] or tertiary [i.e regional or referral facilities]), availability of specialized clinic/ward on site with dedicated staff for TB patients (on site, off site, or not available), physical proximity of HIV/TB clinical services (same facility or same day appointments, cross referral between HIV and TB service points, provision of TB and HIV services under the same roof, or none of these models), and active screening for TB performed for all PLWH at enrollment (only symptom screening, symptom screening plus additional diagnostics, or in case of clinical suspicion). Site-level variables were applied to each patient within the site as an individual characteristic. The main outcome variable was death. Patients were followed from TB treatment initiation until death or censoring within the 12 months post-treatment initiation or June 1, 2015. All the non-death events within the 12 months of follow-up were considered censored events.

### Data analysis

Patient characteristics were summarized overall and by each bacteriologic test group. Categorical variables were summarized using frequencies and proportions; continuous variables were summarized using medians and interquartile ranges. Differences between bacteriologic test groups were assessed using one-way ANOVA or Kruskal-Wallis tests for continuous variables and Pearson Chi-squared test or Fisher’s exact tests for categorical variables. Cumulative incidence of survival, stratified by bacteriologic test group, was estimated using the Kaplan-Meier method. The log-rank test was used for testing equality of survival among the levels of categorical covariates. The Cox proportional hazard model was used to estimate the univariate and multivariate associations of covariates and the hazard for death. Independent variables included in this model were sex, age, BMI, CD4 count, ART status at TB treatment initiation, TB disease site, IeDEA region, and facility level. These variables were selected a priori because of their associations with adverse TB treatment outcomes in prior studies [21, 36–44]. The other site variables were not included in the model due to co-linearity with each other and the facility level variable.

The proportional hazards assumption was assessed using the supremum-type goodness of fit test. The impact of missingness on the observed hazard associations was assessed by refitting the models after imputing missing covariate values. In this analysis, missing values for CD4 count, BMI, and ART status at TB treatment initiation were multiply imputed using the fully conditional specification (FCS), where these covariates were assumed to be jointly distributed [45]. We did not impute missing values for cases with missing facility level, TB disease site, or simultaneously missing BMI, ART status, and CD4 count. Imputation followed Rubin’s scheme: missing values were imputed 100 times; Cox proportional hazards models were fit for the 100 complete datasets; and, results were pooled to obtain overall effect estimates [46, 47]. Imputation analysis was performed using the PROC MI procedure in SAS [48]. Analyses were performed at the 0.05 alpha level.

In a secondary analysis comparing the hazard for death among those with any bacteriologic test (positive or negative) vs. no test results, we used logistic regression to build a propensity score model for the log odds of having any bacteriologic test given imbalance of covariates in the test groups (i.e. any test vs. no test). The independent variables included in this model were the same as those used in the primary analysis. Missing values for CD4 count, BMI, and ART status were also multiply imputed. The imputation and propensity model were performed as follows: 1) missing values were imputed separately for those with and without a bacteriologic test; 2) propensity model was fit; 3) stabilized inverse probability weights (IPWs) were constructed using the predicted propensities, checking if the mean of weights was close to one (S3 Table); 4) the proportional hazards model was fit for time to death conditional on having any bacteriologic test, weighted by the IPWs to obtain an estimate of log hazards ratio and its robust standard error estimate. These steps were repeated 100 times and the resulting log-hazards ratio estimates and their standard errors pooled using Rubin’s rules.

## Results

### Patient and site-level characteristics

Among 2,140 patients in the database, a total of 49 were excluded for initiating TB treatment outside of the study period (n=32) or before the TB diagnosis date (n=1), documentation errors (n=13), and receiving TB treatment regimens for drug-resistant TB (n=3). Thus, 2,091 patients were included in the analysis. These patients received care in 12 countries in the four participating IeDEA regions (Fig 1).

**Fig 1.**
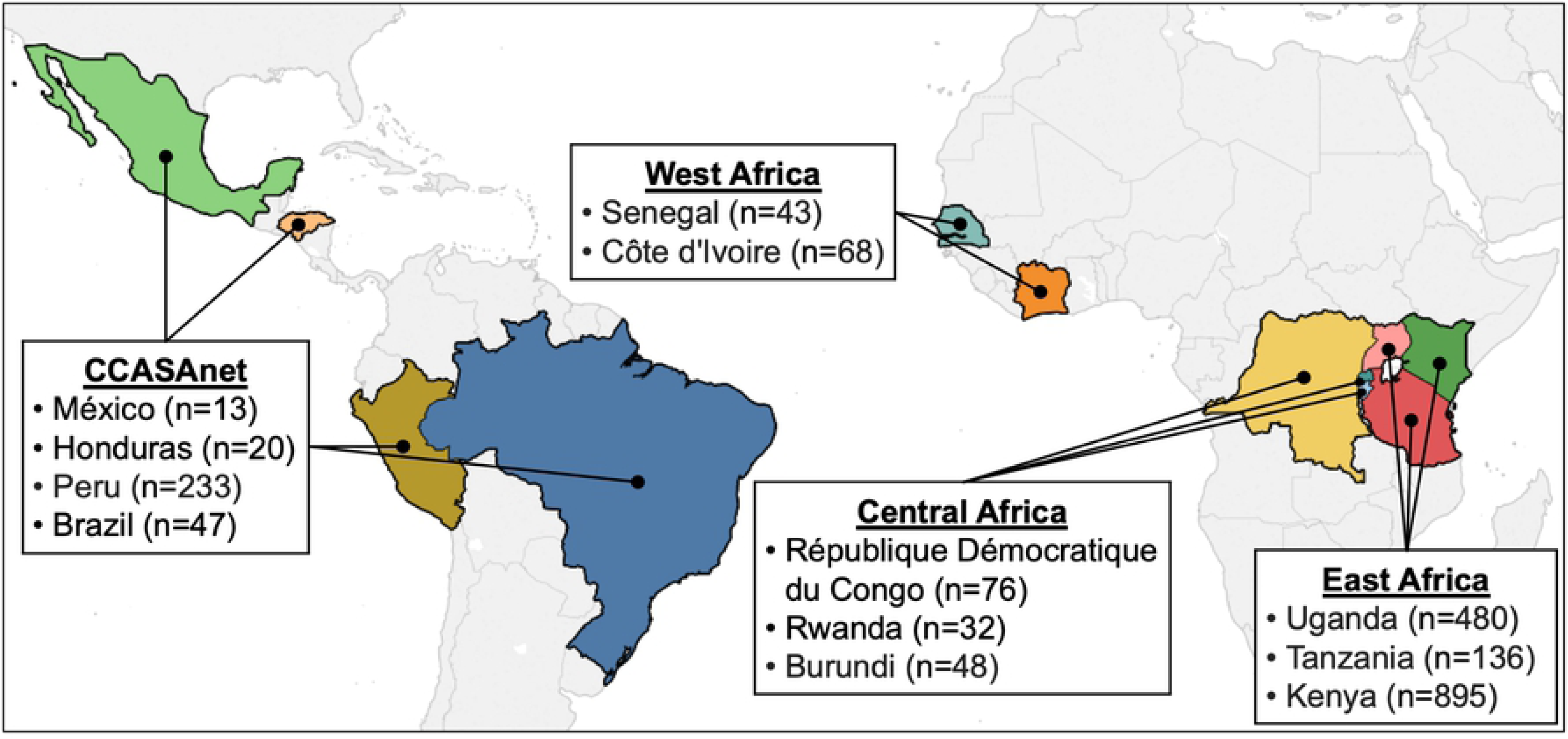
Patients included in the analysis by IeDEA region and country. Numbers in parentheses indicate the number of patients contributed by participating IeDEA programs in each country. Map created in January 2019 by John Humphrey using Tableau Public 2018.3.2 (Tableau Software, Seattle, WA).

A total of 615 (29%) had positive bacteriologic tests for TB, 907 (43%) had negative tests, and 569 (27%) had no test results (Table 1). The median age was 36 years, 44% were female, and the median BMI was 19 kg/m^2^. Overall, 52% were on ART at TB treatment initiation, and the proportions of patients in each bacteriologic test group who were on ART at TB treatment initiation were: positive test (56%), negative test (52%), and no TB test results (38%) (*P* = 0.069). The CD4 count was ≤ 200 cells/mm^3^ in 53% of patients. A total of 79% had pulmonary TB and 20% had extrapulmonary TB; the TB disease site was not specified in 1% of patients. The proportions of patients with extrapulmonary TB were significantly different between the three bacteriologic groups, with extrapulmonary TB in 35% of patients with no TB test result, compared to 8% of patients with a positive test and 19% with a negative test (*P* < 0.001). The most commonly reported extrapulmonary sites were bone and joint (34%) and pleura (22%).

**Table 1.** Patient characteristics stratified by TB bacteriologic test status.

| Characteristic | Bacteriologic Test Status |  |  |  | P value<sup>a</sup> |
| --- | --- | --- | --- | --- | --- |
|  | Total<br>*N* = 2091<br>n (%) | Positive test<br>*N* = 615<br>n (%) | Negative test<br>*N* = 907<br>n (%) | No test result<br>*N* = 569<br>n (%) |  |
| Age, median years (IQR) | 36 (30-43) | 35 (30-42) | 36 (30-43) | 36 (29-44) | 0.779 |
| Female sex | 910 (44) | 266 (43) | 385 (42) | 259 (46) | 0.505 |
| BMI, median mg/kg<sup>2</sup> (IQR) | 19 (17-21) | 19 (17-21) | 19 (17-21) | 20 (17-22) | 0.001 |
| Missing | 374 (18) | 86 (14) | 89 (10) | 199 (35) |  |
| ART status at TB treatment initiation |  |  |  |  | 0.069 |
| On ART | 1084 (52) | 346 (56) | 475 (52) | 216 (38) |  |
| Not on ART | 790 (38) | 215 (35) | 359 (40) | 263 (46) |  |
| Missing | 217 (10) | 54 (9) | 73 (8) | 90 (16) |  |
| CD4 count, cells/mm<sup>3</sup> |  |  |  |  | 0.612 |
| < 100 | 716 (34) | 204 (33) | 309 (34) | 203 (36) |  |
| 100-200 | 391 (19) | 114 (19) | 174 (19) | 103 (18) |  |
| 201-350 | 306 (14) | 85 (14) | 148 (16) | 73 (13) |  |
| 351-500 | 161 (8) | 44 (7) | 73 (8) | 44 (8) |  |
| > 500 | 118 (6) | 36 (6) | 59 (7) | 23 (4) |  |
| Missing | 399 (19) | 132 (21) | 144 (16) | 123 (21) |  |
| TB disease site<sup>b</sup> |  |  |  |  | <0.001 |
| Pulmonary | 1646 (79) | 564 (92) | 732 (81) | 350 (62) |  |
| Miliary | 18 (1) | 0 (0) | 13 (2) | 5 (1) | 0.011 |
| Extrapulmonary | 422 (20) | 49 (8) | 172 (19) | 201 (35) |  |
| Pleural | 79 (22) | 3 (10) | 36 (23) | 40 (20) | <0.001 |
| Lymphatic | 73 (20) | 20 (67) | 26 (17) | 27 (13) | 0.149 |
| Bone and/or joint | 12 (34) | 1 (3) | 5 (3) | 6 (3) | 0.129 |
| Abdominal<sup>c</sup> | 34 (10) | 3 (10) | 18 (11) | 13 (7) | 0.027 |
| Pericardial | 7 (2) | 0 (0) | 4 (3) | 3 (2) | 0.276 |
| Genitourinary | 1 (<1) | 1 (3) | 0 (0) | 0 (0) | 0.566 |
| CNS and/or meningeal | 21 (6) | 1 (3) | 13 (8) | 7 (3) | 0.042 |
| Laryngeal | 1 (<1) | 0 (0) | 1 (1) | 0 (0) | 1.000 |
| Other | 25 (7) | 0 (0) | 11 (7) | 14 (7) | <0.001 |
| Not specified | 23 (1) | 2 (<1) | 3 (<1) | 18 (3) | <0.001 |

ART, antiretroviral treatment; BMI, body mass index; CNS, central nervous system; IQR, interquartile range; TB, tuberculosis
<sup>a</sup> P values comparing the three groups were calculated using ANOVA F-test, chi-square test, Kruskal-Wallis test, and Fisher's exact test.
<sup>b</sup> Percentages of extrapulmonary sites refer to the total extrapulmonary sites; each patient could have > 1 extrapulmonary site.
<sup>c</sup> Includes peritoneum, omentum, liver, spleen, and colon.

The East Africa IeDEA region contributed 72% of patients overall (Table 2). In this region, twice as many patients had a negative bacteriologic test (n=734) compared to a positive test (n=397) or no test (n=380). In contrast, more than twice as many patients in West Africa had either a positive (n=40) or negative test (n=52) compared to no test (n=19). Overall, more patients attended facilities that were peri-urban (47%) than urban (26%) or rural (23%). Among rural facilities, more patients had a negative test (n=324) than either a positive test (n=84) or no test results (72). A total of 88% of patients overall attended tertiary facilities, and there were more patients attending these facilities in the group with negative tests (n=812) than the groups with positive tests (n=472) or no test results (n=485). Finally, 93% of patients attended facilities with a specialized TB clinic/ward available on site, 69% attended facilities with same facility or same day HIV/TB service appointments, and 81% attended facilities with active screening for TB among all PLWH at enrollment that included symptom screening plus additional diagnostics. Site-level differences between the three test groups were significant for all characteristics measured (*P* < 0.001 for all).

**Table 2.** Patient distribution by site characteristics, stratified by TB bacteriologic test status.

| Characteristic | Bacteriologic Test Status |  |  |  | P value<sup>a</sup> |
| --- | --- | --- | --- | --- | --- |
|  | Total<br>N = 2091<br>n (%) | Positive test<br>N = 615<br>n (%) | Negative test<br>N = 907<br>n (%) | No test<br>N = 569<br>n (%) |  |
| **IeDEA region** |  |  |  |  | <0.001 |
| CCASAnet | 313 (15) | 126 (21) | 77 (8) | 110 (19) |  |
| Central Africa | 156 (8) | 52 (8) | 44 (5) | 60 (11) |  |
| East Africa | 1511 (72) | 397 (64) | 734 (81) | 380 (67) |  |
| West Africa | 111 (5) | 40 (7) | 52 (6) | 19 (3) |  |
| **Setting** |  |  |  |  | <0.001 |
| Urban | 537 (26) | 198 (32) | 151 (17) | 188 (33) |  |
| Peri-urban | 986 (47) | 286 (46) | 393 (43) | 307 (54) |  |
| Rural | 480 (23) | 84 (14) | 324 (36) | 72 (13) |  |
| Missing | 88 (4) | 47 (8) | 39 (4) | 2 (<1) |  |
| **Facility level** |  |  |  |  | <0.001 |
| Secondary | 234 (12) | 96 (15) | 56 (6) | 82 (14) |  |
| Tertiary | 1769 (88) | 472 (77) | 812 (90) | 485 (85) |  |
| Missing | 88 (4) | 47 (8) | 39 (4) | 2 (<1) |  |
| **Specialized TB clinic/ward on site** |  |  |  |  | <0.001 |
| Yes, on site | 1951 (93) | 550 (89) | 858 (95) | 543 (95) |  |
| Yes, off site / by referral | 20 (1) | 3 (<1) | 5 (<1) | 12 (2) |  |
| Not available | 32 (2) | 15 (2) | 5 (<1) | 12 (2) |  |
| Missing | 88 (4) | 47 (8) | 39 (4) | 2 (<1) |  |
| **Physical proximity of HIV/TB services** |  |  |  |  | <0.001 |
| Same facility or same day appointments | 1440 (69) | 341 (55) | 709 (78) | 390 (69) |  |
| Cross referral between HIV and TB service points | 401 (19) | 158 (26) | 84 (9) | 159 (28) |  |
| Provision of TB and HIV services under the same roof | 114 (5) | 52 (8) | 44 (5) | 18 (3) |  |
| None of these models | 48 (2) | 17 (3) | 31 (3) | 0 (0) |  |
| Missing | 88 (4) | 47 (8) | 39 (4) | 2 (<1) |  |
| **Active screening for TB for all PLWH at enrollment** |  |  |  |  | <0.001 |
| All, but only symptom screening | 199 (10) | 91 (15) | 32 (4) | 76 (13) |  |
| All, symptom screening plus additional diagnostics<sup>b</sup> | 1689 (81) | 425 (69) | 785 (86) | 479 (84) |  |
| In case of clinical suspicion | 115 (5) | 52 (8) | 51 (6) | 12 (2) |  |
| Missing | 88 (4) | 47 (8) | 39 (4) | 2 (<1) |  |

CCASAnet, Caribbean, Central and South America; PLWH, people living with HIV; TB, tuberculosis
<sup>a</sup> ANOVA F-test, chi-square test, Kruskal-Wallis test, Fisher's exact test performed for each variable as appropriate.
<sup>b</sup> Additional testing examples include sputum AFB, sputum induction, gastric lavage, tissue biopsy, chest x-ray, gene x-pert, urine lipoarabinomannan (LAM), tuberculin skin testing.

AFB smear was performed in 71% of patients (1,493 of 2,091) (Table 3). Culture and NAAT were performed in 8% and 3% of patients, respectively. Among 618 patients with a positive test (whereby ≥ 1 test type may be performed for each patient), 90% had a positive AFB smear, 16% had a positive culture, and 6% had a positive NAAT. Among 907 patients with a negative test (also including ≥ 1 test type), 99% had a negative AFB smear, 6% had a negative culture, and 3% had a negative NAAT. Among patients who had results from more than one test type reported, 75% with a positive AFB smear also had a positive culture (64 of 85) and 35% with a negative AFB smear also had a positive culture (27 of 78) (S4 Table). Among those who had AFB smear and NAAT results reported, 63% with a positive smear had a positive NAAT (10 of 16) and 38% with a negative smear had a positive NAAT (14 of 37).

**Table 3.** Summary of bacteriologic test results.

| Characteristic | Bacteriologic Test Status |  |  |
| --- | --- | --- | --- |
|  | Total tests<br>*N* = 2091<br>n (%) | Positive test<br>*N* = 615<br>n (%) | Negative test<br>*N* = 907<br>n (%) |
| **AFB smear** |  |  |  |
| Positive | 553 (26) | 553 (90) | 0 (0) |
| Negative | 940 (45) | 41 (7) | 899 (99) |
| Not performed / no result | 598 (29) | 21 (3) | 8 (1) |
| **Culture** |  |  |  |
| Positive | 98 (5) | 98 (16) | 0 (0) |
| Negative | 75 (4) | 22 (4) | 53 (6) |
| Not performed / no result | 1918 (91) | 495 (80) | 854 (94) |
| **Nucleic acid amplification** |  |  |  |
| Positive | 38 (2) | 38 (6) | 0 (0) |
| Negative | 34 (2) | 11 (2) | 23 (3) |
| Not performed / no result | 2019 (96) | 566 (92) | 884 (97) |

AFB, acid-fast bacilli

### Mortality outcome

A total of 243 (12%) deaths were reported in the study cohort: 64 (10%) deaths among those with a positive test, 99 (11%) among those with a negative test, and 80 (14%) in those with no test results (*P* = 0.099). The ordering of the survival function in the Kaplan-Meier curve suggested a significant difference in the cumulative incidence of survival between the three bacteriologic test groups. (*P* = 0.017) (Fig 2).

**Fig 2.**
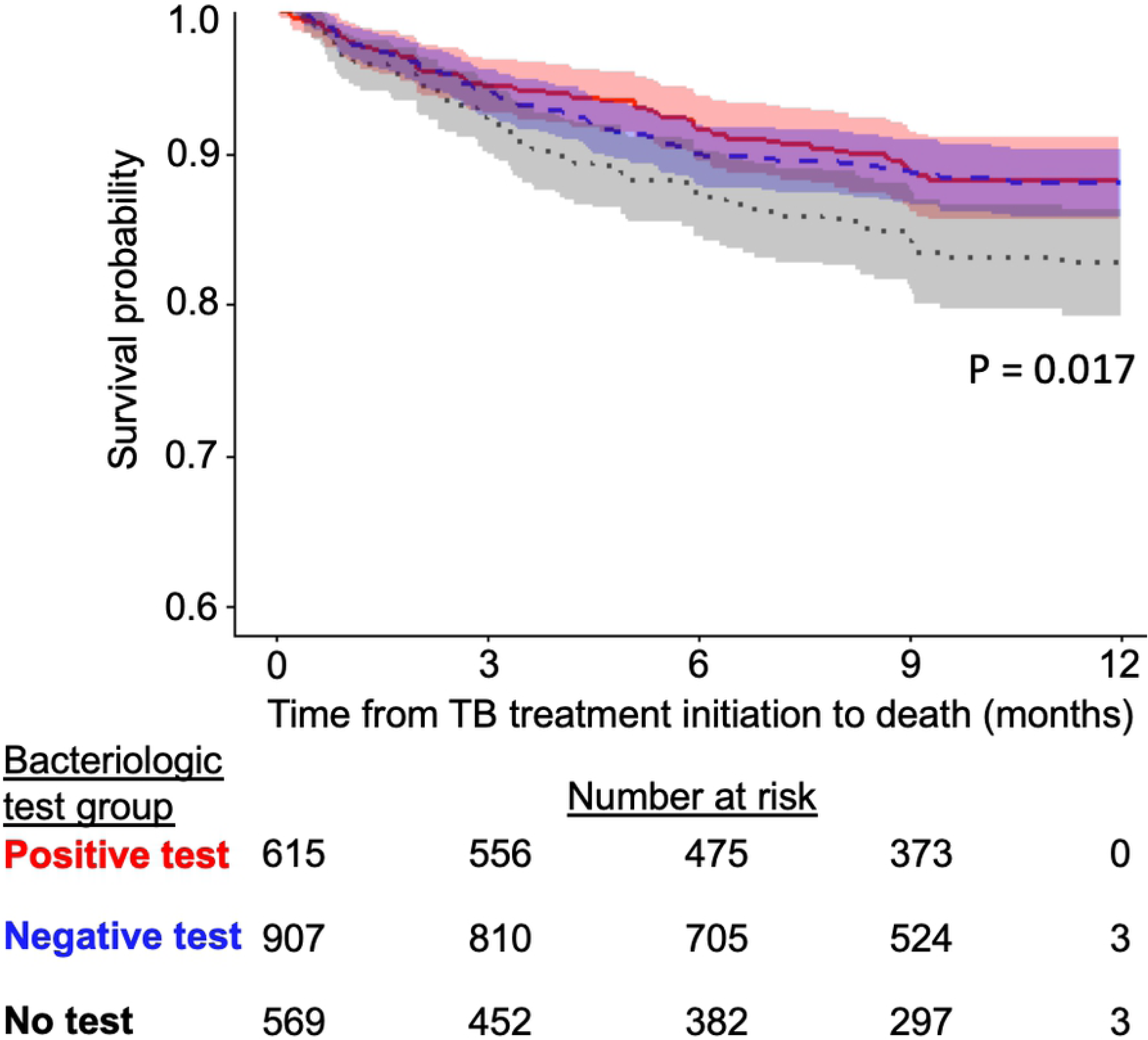
Cumulative incidence of survival after TB treatment initiation, stratified by TB bacteriologic test status. Red, positive test group; blue, negative test group; grey, no test result group.

### Complete case analyses

In the unadjusted and adjusted complete case analyses (adjusted for age, sex, BMI, ART status at TB treatment initiation, CD4 count, TB disease site, region, and facility level), the hazards of death were not significantly higher for those treated for TB with no TB bacteriologic test results (adjusted HR [aHR] 0.87, 95% CI 0.49-1.55) or negative TB tests (aHR 1.19, 95% CI 0.73-1.93) compared to those with positive tests (Table 4). Factors significantly associated with a lower hazard for death in the unadjusted and adjusted analyses included being on ART at TB treatment initiation (aHR 0.57, 95% CI 0.39-0.84) and having a higher CD4 count at TB treatment initiation (e.g. compared to those with CD4 count < 100 cells/mm^3^, the aHR for death among those with a CD4 count of 100-200 cells/mm^3^ was 0.39, 95% CI 0.23-0.65).

**Table 4.** Hazard ratios for death within 12 months following TB treatment initiation.

| Characteristic | Complete Case Analysis<sup>a</sup> |  | Multiple Imputation Analysis<sup>b</sup> |  |
| --- | --- | --- | --- | --- |
|  | Unadjusted HR<br>*N* = 1258 | Adjusted HR<sup>c</sup><br>*N* = 1258 | Unadjusted HR<br>*N* = 1953 | Adjusted HR<sup>c</sup><br>*N* = 1953 |
|  | HR (95% CI) | HR (95% CI) | HR (95% CI) | HR (95% CI) |
| **Bacteriologic Test** |  |  |  |  |
| Positive | 1 (Ref.) | 1 (Ref.) | 1 (Ref.) | 1 (Ref.) |
| Negative | 1.34 (0.86-2.09) | 1.19 (0.73-1.93) | 1.18 (0.85-1.64) | 1.28 (0.91-1.81) |
| None | 1.21 (0.70-2.08) | 0.87 (0.49-1.55) | 1.57 (1.11-2.23) | 1.56 (1.08-2.26) |
| **Age (years)** | 1.01 (0.99-1.03) | 1.01 (0.99-1.02) | 1.02 (1.004-1.03) | 1.02 (1.003-1.03) |
| **Sex** |  |  |  |  |
| Female | 1 (Ref.) | 1 (Ref.) | 1 (Ref.) | 1 (Ref.) |
| Male | 1.23 (0.85-1.78) | 1.15 (0.78-1.69) | 1.24 (0.95-1.63) | 1.20 (0.91-1.59) |
| **BMI (mg/kg<sup>2</sup>)** | 0.97 (0.92-1.02) | 0.98 (0.92-1.03) | 0.97 (0.93-1.01) | 0.97 (0.93-1.002) |
| **ART status at TB treatment initiation** |  |  |  |  |
| Not on ART | 1 (Ref.) | 1 (Ref.) | 1 (Ref.) | 1 (Ref.) |
| On ART | 0.63 (0.44-0.91) | 0.57 (0.39-0.84) | 0.64 (0.48-0.85) | 0.61 (0.47-0.80) |
| **CD4 count** |  |  |  |  |
| < 100 | 1 (Ref.) | 1 (Ref.) | 1 (Ref.) | 1 (Ref.) |
| 100-200 | 0.39 (0.23-0.66) | 0.39 (0.23-0.65) | 0.66 (0.46-0.95) | 0.69 (0.49-0.97) |
| 201-350 | 0.44 (0.26-0.75) | 0.42 (0.24-0.71) | 0.56 (0.37-0.85) | 0.56 (0.38-0.81) |
| 351-500 | 0.16 (0.05-0.51) | 0.15 (0.05-0.48) | 0.47 (0.25-0.87) | 0.46 (0.27-0.79) |
| > 500 | 0.21 (0.07-0.66) | 0.19 (0.06-0.62) | 0.26 (0.10-0.68) | 0.27 (0.12-0.64) |
| **TB disease site** |  |  |  |  |
| Pulmonary | 1 (Ref.) | 1 (Ref.) | 1 (Ref.) | 1 (Ref.) |
| Extrapulmonary | 1.10 (0.70-1.72) | 1.10 (0.68-1.76) | 1.21 (0.88-1.64) | 1.09 (0.79-1.51) |
| **IeDEA Region** |  |  |  |  |
| CCASAnet | 1 (Ref.) | 1 (Ref.) | 1 (Ref.) | 1 (Ref.) |
| Central Africa | 1.04 (0.36-3.01) | 1.08 (0.35-3.33) | 0.92 (0.53-1.62) | 0.52 (0.27-1.02) |
| East Africa | 0.97 (0.52-1.81) | 0.98 (0.49-1.96) | 0.81 (0.58-1.13) | 0.77 (0.54-1.11) |
| West Africa | 1.51 (0.62-3.64) | 0.89 (0.25-3.22) | 1.06 (0.53-2.10) | 0.38 (0.16-0.91) |
| **Facility level** |  |  |  |  |
| Secondary | 1 (Ref.) | 1 (Ref.) | 1 (Ref.) | 1 (Ref.) |
| Tertiary | 0.69 (0.40-1.18) | 0.63 (0.27-1.47) | 0.61 (0.43-0.88) | 0.41 (0.25-0.67) |

ART, antiretroviral treatment; BMI, body mass index; CCASAnet, Caribbean, Central and South America; HR, hazard ratio; CI, confidence interval; Ref., reference; TB, tuberculosis
<sup>a</sup> Includes all patients without missing values for the variables listed in the table.
<sup>b</sup> Missing values were imputed for CD4 count, BMI, and ART status at TB treatment initiation. This model includes all patients without missing values for the variables listed in the table, including those for whom CD4, BMI, and ART status was imputed.
<sup>c</sup> Adjusted for all of the variables listed in the table.

### Multiple imputation analyses

Of 2,091 patients in the study, 1,258 (60%) were complete cases with respect to the analysis model. Of the 833 subjects with missing values, we did not perform imputation for 138 cases. Thus, each imputed dataset had 1,953 subjects. After imputation of missing CD4 count, BMI, and ART status at TB treatment initiation, the unadjusted multiple imputation analysis showed that the hazard for death was significantly higher among those with no test result compared to those with a positive test (HR 1.57, 95% CI 1.11-2.23) (Table 4). In this analysis, older age was significantly associated with a higher hazard for death (HR 1.02, 95% CI 1.004-1.03). Factors significantly associated with a lower hazard for death included being on ART at TB treatment initiation (HR 0.64, 95% CI 0.48-0.85), having a higher CD4 count (e.g. HR for those with CD4 count of 100-200 cells/mm^3^ was 0.66, 95% CI 0.46-0.95), and tertiary facility level (HR 0.61, 95% CI 0.43-0.88). In the adjusted analysis (adjusted the same covariates as in the complete case analysis), the hazard for death was significantly associated with having no test result compared to a positive test (aHR 1.56, 95% CI 1.08-2.26), and there was no difference between those with a positive test and negative test (aHR 1.28, 95% CI 0.91-1.81). Similar to the unadjusted analysis, factors associated with a lower hazard for death included being on ART at TB treatment initiation, having a higher CD4 count and tertiary facility level, as well as West Africa IeDEA region (aHR 0.38, 95% CI 0.16-0.91).

### Secondary analysis

The propensity model for the log-odds of having any bacteriologic test (positive or negative) vs. no test result demonstrated that CD4 count, BMI, TB disease site, facility level, and IeDEA region were each associated with the log odds of having a test (S5 Table). In the odds ratio scale, the adjusted odds of having any test among those with extrapulmonary TB, for example, was 45% lower than the odds for those with pulmonary TB (*P* < 0.0001) (S6 Table). This model was used to estimate the propensity scores, which were in turn used to compute the stabilized IPWs. For each of the 100 analyses combining multiple imputation and propensity score analysis, the stabilized IPWs were found to have an average close to 1. The stabilized IPWs were then used in fitting 100 adjusted Cox models for time to death conditional on whether or not a bacteriologic test had been performed (i.e. test vs. no test). The pooled results for the IPW proportional hazards models demonstrated that the adjusted hazard for death was 28% lower among those with any bacteriologic test compared to those with no test results, but the effect was not significant at the 0.05 alpha level (aHR 0.81, 95% CI 0.61-1.07) (S7 Table).

## Discussion

In this study, PLWH treated for TB in our cohorts in the absence of TB bacteriologic test results had a higher adjusted hazard for death than those treated for TB with positive TB bacteriologic test results. Unlike those with bacteriologically confirmed TB disease (i.e. the positive test group), it is plausible that those with no bacteriologic test results were a heterogenous population of individuals with TB as well as other life-threatening diseases (e.g. opportunistic infections, cancers, chronic lung diseases) that may have mimicked TB but advanced untreated while the patient received TB treatment, resulting in excess mortality. This, along with the 12-month mortality outcome, suggests that not all patients who initiated TB treatment had TB disease and not all deaths were TB-attributable.

Differences in TB disease site or severity between those with positive and no test results may also have influenced mortality. As expected, the proportion of patients with pulmonary TB was higher among those with positive tests (92%) compared to those with no test results (62%) (*P* < 0.001). This reflects the challenge of acquiring a bacteriologic diagnosis in suspected extrapulmonary TB (which often requires invasive tissue sampling) compared to pulmonary TB (which requires sputum sampling) in resource-constrained settings. Additionally, bacteriologic testing for extrapulmonary TB is less sensitive in general compared to pulmonary TB, extrapulmonary TB is more common in PLWH, certain extrapulmonary TB types (e.g. disseminated or meningitis) are associated with especially high mortality in PLWH, and extrapulmonary TB has been associated with delays in diagnosis [40, 42, 43, 49–52]. These features may also have influenced clinicians to forego bacteriologic testing and initiate empiric TB treatment among individuals that were already at increased risk of death. TB disease site was not associated with mortality in our analyses, but we could not account for group variability in extrapulmonary TB sites that may have influenced this outcome.

We also found that PLWH attending tertiary facilities had a lower adjusted hazard for death compared to those attending secondary facilities. Compared to tertiary facilities, secondary facilities may have had less capacity to evaluate and manage TB, or its alternative diagnoses, in PLWH [53]. This finding may also have been biased by differences in the completeness of death ascertainment between higher and lower-resourced facilities, the latter being potentially more susceptible to undocumented loss to follow-up or transfer events not captured in our dataset [54]. Differences in vital status ascertainment may also have accounted for the regional differences in the hazards for death (West Africa vs. CCASAnet) identified in our study.

Consistent with the literature, we found that being on ART and having higher CD4 count at TB treatment initiation were both strongly protective against mortality. This underscores the fundamental importance of early ART initiation and immune preservation on survival, regardless of the presence or result of TB bacteriologic testing, in resource-constrained settings. Advanced HIV immunosuppression is known to be a critical risk factor for TB-related mortality (including mortality ≥ 1 year after completion of TB therapy), and the scale-up of ART coverage has been associated with marked reductions in TB incidence and mortality in countries with high HIV and TB burdens [20, 21, 40, 41, 55–57].

We did not find a significant difference in the adjusted hazard for death between those treated for TB with any bacteriologic test (positive or negative) versus no test results in the secondary analysis. This finding is consistent with a study in Brazil which found no difference in mortality risk between PLWH who did or did not undergo TB bacteriologic testing [17]. This argues against the notion that the presence of TB bacteriologic testing is a marker of better-resourced sites (and therefore reduced mortality), as patients with a bacteriologic test would have been expected to have a lower mortality than those who were not tested if that were the case. It is possible that patients in both groups had similar survival simply by being engaged in facility-based care.

Finally, we found no significant difference in the hazard for death between those with positive or negative TB bacteriologic tests (Fig 2 and Table 3). This lack of significance may in part be related to limited statistical power, as the adjusted hazard for death in those with negative tests was similar to that for those with no test results (aHR 1.28 vs. 1.56). Nevertheless, this finding contrasts with prior studies finding that smear-negative TB is associated with increased mortality compared to smear-positive TB in PLWH. This finding has been attributed to paucibacillary disease in the setting of advanced HIV, delay in diagnosis, and other opportunistic and non-communicable diseases [14, 21–23, 58–65]. Similar findings have been shown with respect to culture results among PLWH treated for TB, but no such studies have been performed for NAAT to our knowledge [66]. The lack of mortality difference in our study could also be related to the combination of smear, culture, and NAAT used to define the groups, overdiagnosis of TB in the negative test group (yielding lower than expected mortality), or higher bacterial burden due to more extensive TB disease or inadequate treatment of drug-resistant TB in the positive test group (yielding higher than expected mortality) [67]. The 12-month mortality outcome used in our study, which was selected due to limitations in the dataset, may also have influenced this finding. A study from the Democratic Republic of Congo, for example, found that patients treated for smear-negative TB had a higher risk of death within two months of TB treatment initiation, but not after, compared to those treated for smear-positive TB [62].

Our study has strengths and limitations. Strengths include its large sample size from diverse global regions and HIV/TB care programs and use of routine program data which likely reflects typical care environments in the study settings. The use of routine program data is also a limitation, as the completeness of vital status ascertainment may have been affected by loss to follow-up or transfer events not captured in our dataset. We cannot be certain that patients with no TB bacteriologic test results documented in their records did not actually have testing performed. Still, the integration of TB-HIV services at the majority of sites supports that TB test results would have been recorded in the medical records and case report forms if they were available. Few patients had culture or NAAT performed, which limits the generalizability of our study in settings where these tests are performed more commonly. We used multiple imputation to reduce bias due to missing data, but the mortality estimate could still be biased if missingness depended not only on the variables we used to impute missing values but also on the missing values themselves (i.e., missing not at random).

In conclusion, PLWH treated for TB with no TB bacteriologic test results in our study were more likely to die than those who were treated and had positive tests. Every effort should be made to establish a diagnosis of TB prior to initiating TB treatment in resource-constrained settings. Further research is needed to understand the causes of death among PLWH treated for TB in the absence of positive bacteriologic test results.

## Acknowledgements

We are grateful for the participation of the following IeDEA Collaborative sites: Brazil – INI-Fiocruz; Burundi – CPAMP-CHUK; Côte d’Ivoire – Centre Intégré de Recherches Biocliniques d’Abidjan (CIRBA), Centre Hospitalier Universitaire de Cocody, CREF/SMIT, Medicine Interne; Honduras - IHHS; Kenya –AMPATH; Mexico - INCMNSZ; Peru – IMTAvH, CoVIHS; République Démocratique du Congo – Bomoi Health Centre, Kalembelembe Paediatric Hospital; Rwanda – Rwanda Military Hospital; Tanzania – National AIDS Control Programme, Tumbi Regional Hospital, National Institute for Medical Research, Mwanza Research Centre-Kisesa Clinic; Uganda – Masaka Regional Hospital. A complete listing of participating programs and members can be found in S8 Table.

## Supporting Information

**S1 Table. STROBE checklist**.

**S2 Table. Independent Ethics Committee or Institutional Review Board approvals obtained by participating IeDEA sites**.

**S3 Table. Computation of stabilized inverse probability weights**.

**S4 Table. Summary of the number (%) of positive and negative bacteriologic test results in patients with more than one test type performed.** (A) AFB smear + culture, (B) AFB smear + NAAT, (C) Culture + NAAT.

**S5 Table. Propensity model results in the log of odds ratio scale**.

**S6 Table. Propensity model results in odds ratio scale**.

**S7 Table. Inverse probability weighted Cox proportional hazards model for hazard of death within 12 months following TB treatment initiation conditional on receiving a TB bacteriologic test (positive or nagative).** Includes imputation of CD4 count, body mass index (BMI), and antiretroviral treatment (ART) status at TB treatment initiation.

**S8 Table: Membership of the International Epidemiology Databases to Evaluate AIDS (IeDEA) iparticipating programs**.

